# Immune response to co-administration of bovine tuberculosis and contraceptive vaccines in badgers (*Meles meles*)

**DOI:** 10.1101/2025.02.12.637632

**Authors:** Kate L. Palphramand, Paul Anderson, Fiona Bellamy, Colin Birch, Dipesh Davé, Doug Eckery, Matt Gomm, Rachel Jinks, Giovanna Massei, Rebecca Pinkham, Sabah Rahou, Hugh Simmons, Flavie Vial, Gareth A. Williams, Graham C. Smith

## Abstract

Controlling bovine tuberculosis (bTB) in livestock is often hindered by the presence of a wildlife reservoir, such as the Eurasian badger in the UK and Ireland. Vaccinating badgers against bTB can reduce the severity of *Mycobacterium bovis* infection and potential onwards transmission to cattle, badgers and other species, thus combined with population control might provide additional benefits (and reference models). To evaluate the effects of co-administration of a bTB vaccine (BCG) and a contraceptive vaccine (GonaCon), captive badgers were injected intramuscularly with either BCG only, GonaCon only or BCG+GonaCon (phase 1). The duration of immune response to BCG vaccination and booster vaccination was also evaluated in the BCG only group over a 24-month period (phase 2). The cellular immune responses to purified protein derivatives PPDA and PPDB, used to measure the effects of BCG, did not differ significantly between the BCG only and BCG+GonaCon badgers but were significantly different between the BCG only and GonaCon only badgers. Anti-GnRH antibody titres, quantified to measure the effect of GonaCon, peaked at the first sampling point for the majority of badgers in the GonaCon only and BCG+GonaCon groups. Throughout phase 1, lasting three months post-vaccination, badgers in both GonaCon treatment groups maintained high anti-GnRH antibody titres (1:128,000 or above) that are associated with infertility in this species, with no significant difference between the two GonaCon groups. The immune responses to BCG vaccination in the BCG only badgers (phase 2) were elevated at the first sampling point, approximately three weeks after initial vaccination and booster vaccination, generally declining thereafter. The results from the current study suggest that co-administration of BCG and GonaCon do not have notable deleterious effects on either of the vaccines and that the protective effect of BCG is enhanced by booster vaccination. The protective effects afforded by both vaccines when given together, in terms of reduced disease burden or fecundity, should be determined by further studies.

## 1. Introduction

Bovine tuberculosis (bTB) is an infectious zoonotic disease caused by *Mycobacterium bovis*. It has a global distribution and poses a risk to the health of livestock, particularly cattle (Blanco et al., 2022). It also causes significant economic impacts and as a result, many endemic countries have implemented long-running programmes for eradicating tuberculosis in livestock, including Great Britain and Ireland, which have had a compulsory test and slaughter programme for cattle since 1950 (Reynolds, 2006). Despite an initial decline to very low levels of tuberculosis in cattle, bTB incidence started to increase by the early 1990’s with the test and slaughter scheme remaining central to the strategy to stop its spread (Reynolds, 2006).

In the UK and Ireland, bTB eradication programmes are currently in place with an estimated annual cost to taxpayers of over £100 million in Great Britain alone (Defra, 2024). Eradication efforts have been hindered, however, by the presence of the disease in wildlife species, in particular the European badger (*Meles meles*), which is a recognised reservoir host (Palmer, 2013). Badger culls have been carried out by the UK government since the early 1970’s, when badgers were first recognised as being involved in the epidemiology of bTB in cattle (Goodchild & Clifton-Hadley, 2006). However, badger culling can result in complex epidemiological outcomes, with both positive and negative effects on the incidence of bTB in cattle (Griffin et al., 2005; Donnelly et al., 2006). Multiple studies have shown that bTB transmission is complicated, unlikely to be driven by a single mechanism and strongly associated with the local environment and host dynamics of the system (van Tonder et al., 2021; Chang et al., 2023). Although previous research has demonstrated an association between *M. bovis* found in sympatric cattle and badger populations, quantifying directional estimates of transmission have been difficult (Crispell et al., 2019). However, recent evidence from a number of extensive studies suggests that, whilst most disease transmission is intra-specific, occurring within cattle herds or within badger populations, inter-species transmission is also evident with the estimated rate of transmission from badger to cattle being higher than is estimated in the opposite direction for some areas (Crispell et al., 2019; van Tonder et al., 2021), while the reverse has been true in others (Rossi et al., 2022).

Vaccination has long been considered as one tool to help reduce bTB infection in badger populations, with the UK government investing approximately £27 million in the research and development of badger vaccines since the mid-1990s (Benton et al., 2020). Consequently, an injectable vaccine (BadgerBCG, hereafter referred to as BCG) was licensed in 2010, as it was shown to reduce the severity and progression of the disease in both captive (Lesellier et al., 2011) and wild badgers (Chambers et al, 2011), and provide herd immunity for cubs (Carter et al., 2012). In a review of Defra’s Strategy for achieving Officially Tuberculosis Free status for England (Defra, 2014), Godfray et al. (2018) proposed moving from lethal to non-lethal control of bTB in badgers, with injectable BCG vaccine considered the only viable non-cull control option available at the time. The most recent TB eradication strategy proposed by the UK government aims to build on the Godfray et al. (2018) review by combining badger vaccination with a number of other measures. These included an acceleration of work to develop a cattle vaccine, a major survey of the badger population (which was last carried out between 2011-2013) and a wildlife surveillance programme to provide an understanding of the prevalence of bTB in the remaining badger populations, which is largely unknown (Defra, 2024).

In parts of the UK, the density of badgers was considered to be higher than anywhere else in Europe (Smith, 2002), with badger numbers prior to recent culling estimated to have increased in England and Wales since the 1980s to approximately 485,000 (Judge et al., 2017). Whilst vaccination has the potential to control bTB in badgers, which in turn could reduce the incidence of the disease in cattle, using this approach alone would not directly reduce the number of badgers, unlike culling. An alternative non-lethal approach for wildlife disease management could be to combine fertility control and disease vaccination. Fertility control has long been considered a humane method for reducing mammal abundance where conflicts exist between wildlife and human interests (Fagerstone et al., 2010; Massei & Cowan, 2014; Massei, 2023). Carroll et al. (2010) modelled the effect of combining a single-dose contraceptive with rabies vaccination in urban dogs; they concluded that using immunocontraception alongside disease vaccination could improve rabies campaigns by reducing the proportion of the population that required treatment or reducing the duration of the campaign. One of the most frequently used immunocontraceptives developed for mammals, is based on the gonadotropin-releasing hormone (GnRH) (Massei, 2023). Immunisation of both male and female animals induces the production of antibodies against the GnRH, which in turn reduces the concentrations of sex hormones and inhibits reproductive function in both sexes (Miller et al., 2008). In wildlife, a frequently studied immunocontraceptive is the injectable GnRH-based vaccine, GonaCon (Massei, 2023), registered in the USA as a contraceptive for white-tailed deer (*Odocoileus virginianus*), feral horses (*Equus caballus*), feral donkeys (*Equus asinus*) and prairie dogs (*Cynomys ludovicianus*). GonaCon has been tested in a range of mammal species with no adverse effects on the behaviour of males or females and can induce infertility for several years after one or two doses (Cowan et al., 2019; Massei, 2023). Cowan et al. (2019) demonstrated that a single-shot injectable immunocontraceptive given to captive badgers inhibited subsequent cub production the following year, with the longer-term effectiveness of GonaCon in female badgers appearing to reflect maintenance of anti-GnRH antibody titres at or above a putative threshold titre of 1:128,000. These authors suggested that combining fertility control and disease vaccination could be a potentially attractive strategy for managing bTB in badgers.

For some wildlife species, vaccination efforts may be restricted to the use of a single-shot vaccine as it is often difficult to recapture or access individuals to administer a booster dose (Buddle et al., 2018). In contrast, badgers are ideal candidates for administering injectable vaccines as they are relatively easy to capture (due to living in conspicuous setts) and will often re-enter traps on subsequent occasions (e.g. Tuyttens, et al., 1999; Chambers et al., 2010; Byrne et al., 2012; Carter et al., 2012), although there will be a small proportion of badgers that never enter traps (Tuyttens et al., 1999). Badgers are routinely trapped during BCG vaccination operations; several sources have reported on the costs of trapping and vaccinating badgers in a high-density area, with the Gloucestershire Wildlife Trust estimating the average cost over a five-year deployment trial of £266/badger vaccinated and the Welsh Government an estimated £706/badger vaccinated (Gloucestershire Wildlife Trust, 2015). Therefore, the cost-effectiveness of BCG vaccination campaigns might be improved by combining them with the concurrent application of fertility control.

The main aim of the current pilot study was to determine the potential interactions and effect on immune responses when BCG and GonaCon vaccines were injected simultaneously in captive badgers (phase 1). In particular, the study aimed to assess whether the use of GonaCon to control fertility in badgers would reduce the immunity against bTB conferred by the BCG vaccine and vice versa. The study also aimed to investigate the duration of immune response to BCG over 12 months followed by a booster dose (phase 2); this is considered indicative of duration of immunity but without the challenge with an infectious agent (i.e. demonstration of protection against *M. bovis*). Although this approach is more refined than with challenge, it can only provide evidence of immunological correlates of protection but would greatly enhance scientific knowledge in this area.

## 2. Materials and methods

### 2.1 Badgers

For phase 1, 32 adult badgers were trapped in autumn 2018 from Northumberland, which has a very low prevalence of bTB in cattle (APHA, 2023), under the terms of a Natural England licence. Badgers were transferred to APHA Weybridge, UK, and housed in the same social groups they were captured from. Badgers were housed in enriched pens with standard husbandry procedures, ad lib food and water and observed daily by trained staff for monitoring and welfare reasons. Badgers were confirmed bTB-free during a quarantine period using a series of badger Interferon Gamma Release Assays (IGRA), each assay one month apart over a three-month period. Additionally, tracheal and rectal swab clinical samples were submitted for *M. bovis* microbiological culture at each time point and an ELISpot was used to confirm IGRA results at the second sampling point.

The 32 badgers were comprised of nine social groups (Table 1); each individual received either BCG only (five females, four males), GonaCon only (four females, four males) or BCG+GonaCon (nine females, four males). Different individuals within the same social groups received different treatments; although badgers were randomly assigned to their treatment groups, this was limited to avoid an uneven allocation of males and females to the GonaCon groups, which could have biased results (females being the main target in fertility control protocols).

**Table 1.** Number of badgers per social group, number of social groups and treatment allocation during phase 1 (total number of badgers n=32).

| No. of badgers per social group | No. of social groups | No. of badgers receiving BCG only* | No. of badgers receiving GonaCon only | No. of badgers receiving BCG+GonaCon |
| --- | --- | --- | --- | --- |
| 4 | 6 | 1 | 1 | 2 |
| 3 | 2 | 1 | 1 | 1 |
| 2 | 1 | 1 | 0 | 1 |

\* Six of the nine badgers in the BCG only group were taken into phase 2.

Phase 2 investigated the duration of immune response and effect of a second dose of BCG on the cohort of badgers from the BCG only group in phase 1. Three of the nine badgers from the original cohort were euthanised after phase 1; the remaining six badgers were transferred to APHA York, UK, and housed in pairs (for social welfare reasons) based on prior behavioural observations and advice from the Named Veterinary Surgeon and Named Animal Care and Welfare Officer. Badgers were housed in enriched pens with standard husbandry procedures, ad lib food and water and observed daily by trained staff for monitoring and welfare reasons.

### 2.2. Blood sampling

Blood was collected under general anaesthesia at a series of time points to measure immune responses (Table 2). Badgers were given an intramuscular anaesthetic cocktail of ketamine, medetomidine and butorphanol (plus hyaluronic acid in phase 2) with top-up of isoflurane if required. Samples were collected by venepuncture of not more than 10% of circulating blood volume on a single occasion, or 15% over a 28-day period for repeated blood sampling; during phase 2, 50ml of blood was taken from each badger. Blood was collected into heparin or serum-separating Vacutainer® tubes. After sampling, badgers were closely monitored until they had fully recovered. During phase 1, blood sampling was scheduled no more frequently than every third week for the first two to three months to measure BCG immune response, followed by a minimum gap of two months to measure GonaCon, which has a slower effect. Due to welfare reasons and Covid-19 restrictions during 2021, the timeline of blood sampling during phase 2 differed to phase 1.

**Table 2.** Timeline of events (blood sampling, vaccination and euthanasia) during phase 1 and 2 for badgers in the three treatment groups (BCG only, GonaCon only, BCG+GonaCon).

| Day from phase 1 | Day from phase 2 | Date | Event |
| --- | --- | --- | --- |
| 0 | - | 10/09/2019 | Baseline bloods + vaccinations (BCG & GonaCon) |
| 22 | - | 01/10/2019 | Bloods |
| 42 | - | 22/10/2019 | Bloods |
| 62 | - | 11/11/2019 | Bloods |
| 90 | - | 09/12/2019 | Bloods; GonaCon only and BCG+GonaCon badgers euthanised |
| 92 | - | 11/12/2019 | BCG only badgers transferred to York |
| 181 | - | 09/03/2020 | Bloods |
| 373 | 0 | 17/09/2020 | Baseline bloods + BCG booster vaccination (start of phase 2) |
| 399 | 26 | 13/10/2020 | Bloods |
| 644 | 271 | 15/06/2021 | Bloods |
| 741 | 368 | 20-22/ 09/2021 | Bloods; BCG only badgers euthanised |

### 2.3. Vaccination

Badgers were vaccinated under general anaesthesia (Table 2). Both vaccines, *Mycobacterium bovis* BCG Danish Strain 1331 (2-8 × 10^6^ CFU BCG, supplied by AJ Vaccines, Denmark) and GonaCon Equine (GnRH 0.032%, imported under VMD permit from SpayFirst, USA), were stored at 2-8°C until required for vaccination. Each badger received 1ml of vaccine, given by intramuscular injection in the hind leg (in the BCG+GonaCon group, 1ml of BCG and 1ml of GonaCon were administered separately in different hind legs). After vaccination, badgers were monitored for any adverse effects, such as swelling or lameness, before being returned to group housing.

### 2.4. Immune responses

#### 2.4.1. Immune responses to BCG

Response to BCG vaccination was measured using a range of cellular assays. After incubating whole bloods with a range of antigens, the plasma was removed and the samples were run on an Interferon Gamma Release Assay (IGRA) as described by Dalley et al. (2008). In addition, whole bloods were separated to produce peripheral blood mononuclear cells (PBMC). These were also stimulated with antigens to provide additional samples for the IGRA and an enzyme-linked immune absorbent spot (ELISpot) assay to measure the quantity of interferon gamma (IFNγ) produced and the number of reactive PBMCs, respectively. Antigens used to stimulate PBMC cultures in duplicate wells were those used for the routine diagnosis of TB in humans and animals (purified-protein derivatives (PPD) made from *M. bovis* (PPDB) and *M. avium* (PPDA) (APHA Weybridge antigens)). PPDA is used to measure background responses to environmental mycobacteria (*M. avium*), while PPDB contains *M. bovis*-specific antigens (Rhodes, 2015). Although immunological responses to BCG are potentially complex, recent studies suggest that response to PPDB is a relatively strong indicator of effective vaccination (Lesellier et al., 2020).

#### 2.4.2. Immune responses to GonaCon

The efficacy of GonaCon was determined by serum assay for GnRH antibodies by ELISA using the method described by Levy et al. (2004). Serum was separated by centrifugation and stored at −20°C if not assayed immediately. Plates were prepared by adding BSA-GnRH antigen (USDA APHIS, USA) to each well before being stored overnight at 4°C. Plates were then washed with phosphate buffered saline (PBS) with 0.05% TWEEN®20 (P3563, Merck, UK) before adding SynBlock blocking buffer (BUF034C, Bio-Rad, UK). Plates were washed with wash buffer as above before being loaded with serial dilutions of OBT1999 buffer (Bio-Rad, UK) and diluted (1:1000) badger serum in OBT1999 buffer. Pre-bleed serum for the individual badgers and a positive control serum (both diluted 1:1000 in OBT1999 buffer) were also added to the plates. Following incubation of the plates at 25°C for one hour, plates were washed with PBS wash buffer and then loaded with diluted 1:4000 CF2 rabbit x badger IgG-HRP monoclonal antibody (APHA, Weybridge) in PBS wash buffer. Colour was developed by 3,3’5,5’-tetramethylbenzidine dihydrochloride (TMB) substrate (PN34021, Fisher, UK) and absorbances analysed to determine the anti-GnRH antibody titre of each sample. Values were corrected to account for OBT1999 dilutions before assigning titre values.

### 2.5. Statistical analysis

#### 2.5.1. Responses to BCG vaccination

For analysis, IGRA and ELISpot responses of PBMCs after stimulation by PPDA and PPDB antigens were log transformed to make their variance more uniform, as described in Lesellier et al. (2020). The IGRA responses were transformed using log_10_(x-n), where n was chosen to be slightly below the smallest value of the data: for PPDA this was 0.044 (n=0.043), PPDB minimum was 0.039 (n=0.038). The ELISpot responses were multiplied by 5 to give SFU/10^6^ cells, then log transformed using log_10_(5x+5), where x is the raw ELISPot data. The net value PPDB-PPDA was also determined to provide a level of immune response which is *M. bovis* specific. Any value over zero is considered an immune response to vaccination whereas any value below zero (where the response of PPDB is less than PPDA) is classed as no response to vaccination. As such, the binary variable was also considered to avoid the difficulties of analysing a continuous variable with a large peak at zero. This difference was taken after the log transformation described above, i.e. log10(PPDB-0.038)-log10(PPDA-0.043).

Mixed random effects models were constructed to determine statistical differences in immune responses due to treatment group and over time. Treatment and day of study were included as fixed effects (excluding baseline time point day 0, which was pre-vaccination) and individual badgers as a random effect to account for repeated measures on the same animal. Day of study was considered as both a continuous and a factor effect. The interaction between treatment and day was also investigated to check whether the effect of vaccination over time was different for the three treatment groups. Models were compared using the Akaike Information Criterion (AIC), such that a smaller AIC suggested a more parsimonious model.

#### 2.5.2. Responses to GonaCon vaccination

Responses to GonaCon vaccination were measured using anti-GnRH antibody titres. These were calculated by comparing blood samples between pre- and post-vaccination for each individual and expressed as the highest dilution, in which the post-vaccination sample had a meaningfully higher absorbance value than the pre-vaccination sample. Titre observations were reported as powers of 2 (1:X,000), i.e. below threshold, 2, 4, 8, 16 up to 8192 (2^13^). For analysis, titres were logarithm transformed to the series of integers 0-13, with zero representing below threshold. The titre observations were analysed in mixed effect models, which included treatment and time as covariates and badger ID as a random effect to take account of the repeated measurements on the same animals. Three models were constructed: model 1 treated the four time points as independent; model 2 included time as a continuous variable with a simple linear relationship with titre; and model 3 included potential interactions between treatment and time. Models were compared using the AIC.

## 3. Results

### 3.1. Phase 1

#### 3.1.1. Treatment effects on BCG response

The mean cellular immune responses of the badgers in the three treatment groups (BCG only, GonaCon only and BCG+GonaCon) as measured by IGRA and ELISpot are shown in Figures 1 and 2. Immune response to both PPDA and PPDB for the BCG only and BCG+GonaCon groups peaked at the first time point after vaccination (day 22) and declined thereafter, although the responses remained above pre-vaccination levels beyond day 90. The profile was similar whether measured by ELISpot or IGRA. The mean immune response to PPDA and PPDB of badgers in the GonaCon only treatment group was flatter than the BCG treatment groups when measured using IGRA. However, there was a marked peak in immune response to both PPDA and PPDB at 42 days when measured using ELISpot. When the immune response was defined by whether the PPDB response was greater than PPDA (Figure 3), the proportion of animals with a response fluctuated slightly over time for the GonaCon only and BCG+GonaCon groups; in contrast, the proportion in the BCG only group generally increased over time (Figure 3).

**Figure 1.**
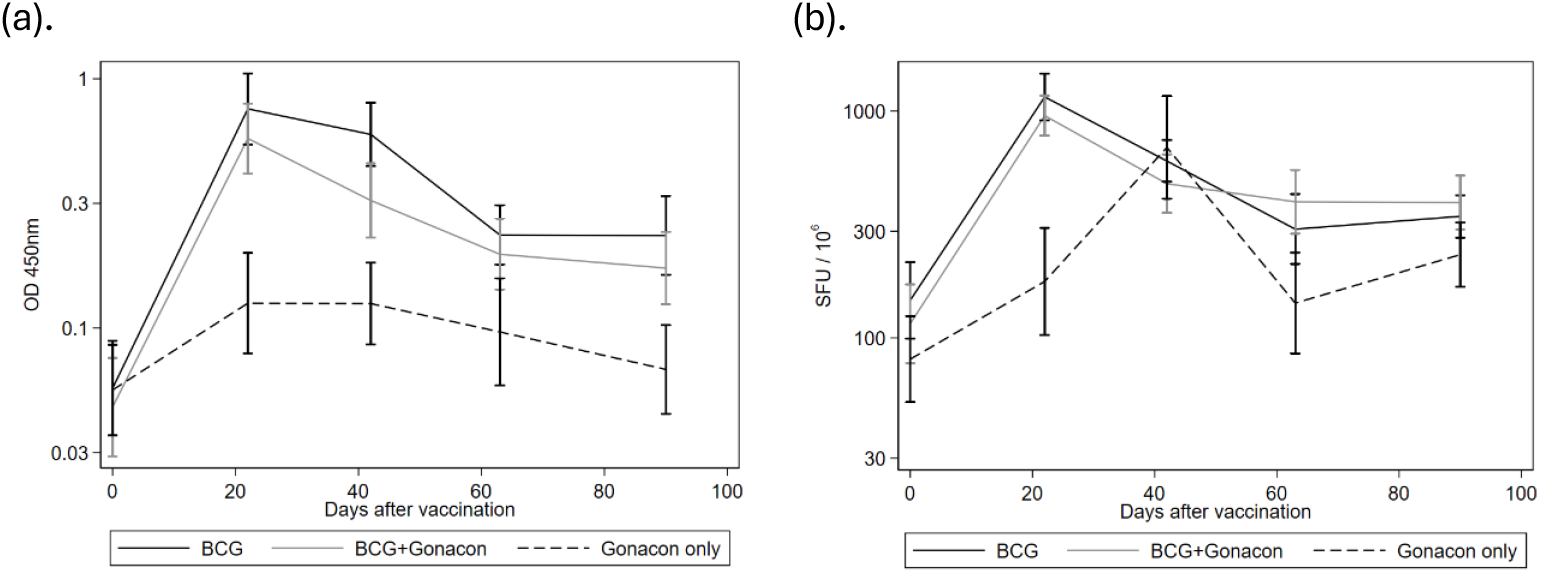
Mean immune response of the three treatment groups (BCG only, n=9, BCG+GonaCon, n=15, GonaCon only, n=8) over the course of phase 1 as measured by (a). IGRA PPDA, (b). ELISpot PPDA (net SFU). Error bars are standard error of the means.

**Figure 2.**
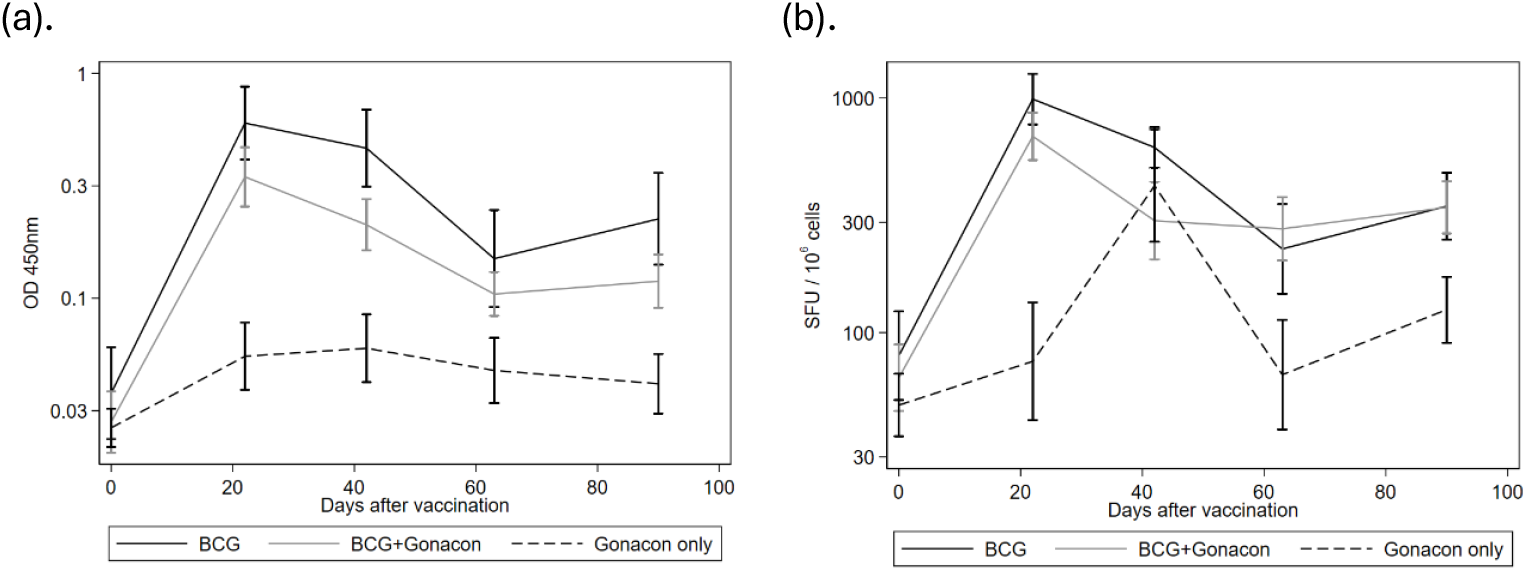
Mean immune response of the three treatment groups (BCG only, n=9, BCG+GonaCon, n=15, GonaCon only, n=8) over the course of phase 1 as measured by (a). IGRA PPDB, (b). ELISpot PPDB (net SFU). Error bars are standard error of the means.

**Figure 3.**
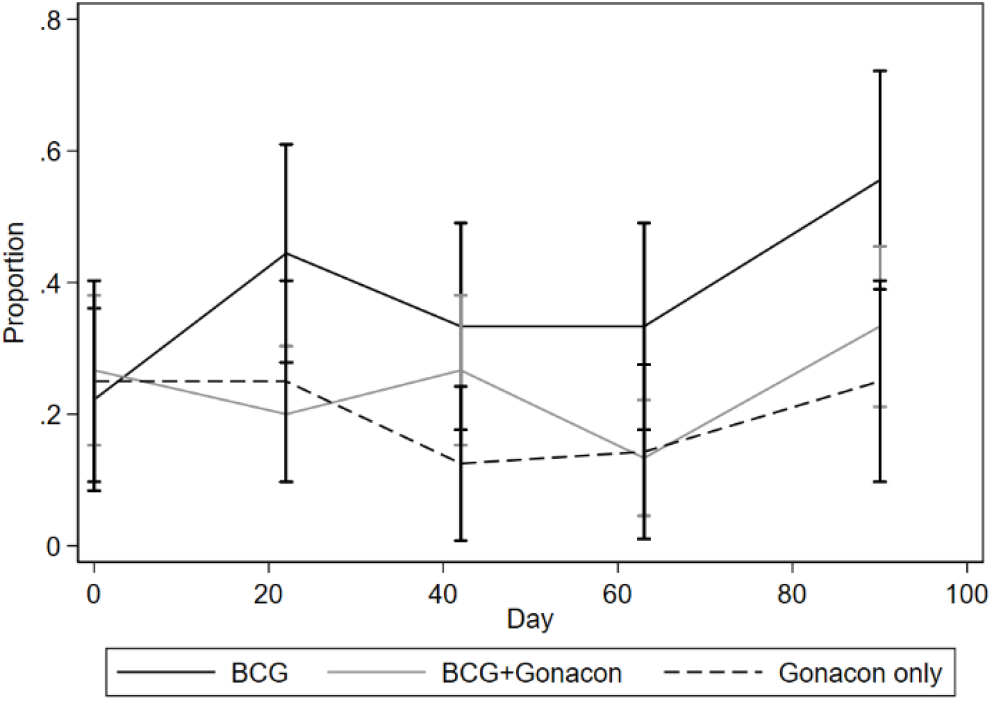
Proportion of animals within the treatment groups (BCG only, n=9, BCG+GonaCon, n=15, GonaCon only, n=8) with IGRA PPDB>PPDA over time. Error bars are standard error of the proportion.

Figures 1 to 3 showed that, in general, animals in the GonaCon only group had the lowest immune response after BCG vaccination, apart from the ELISpot peak at day 42. Five mixed effect models were compared for differences in immune response as measured by IGRA (Table 3) and ELISpot (Table 4). Day of study was found to be best modelled as a factor effect (lower AIC) for all five. The interaction terms treatment:day did not improve the models using IGRA data, but were required to model the ELISpot data (AIC=173 with interactions vs 178 without for PPDA response, and AIC=199 vs 206 for PPDB response).

**Table 3.** Differences in immune response to log-transformed IGRA PPDA and PPDB for (i) BCG+GonaCon compared to BCG only and (ii) GonaCon only compared to BCG only during phase 1.

| Model term | PPDA |  |  | PPDB |  |  |
| --- | --- | --- | --- | --- | --- | --- |
|  | Coefficient | SE | P value | Coefficient | SE | P value |
| BCG+GonaCon* | -0.148 | 0.180 | 0.410 | -0.254 | 0.162 | 0.118 |
| GonaCon* | -0.610 | 0.207 | 0.003 | -0.801 | 0.187 | <0.001 |
| Day 42† | -0.146 | 0.072 | 0.042 | -0.122 | 0.072 | 0.092 |
| Day 63† | -0.403 | 0.072 | <0.001 | -0.441 | 0.073 | <0.001 |
| Day 90† | -0.453 | 0.072 | <0.001 | -0.368 | 0.072 | <0.001 |

\* vs BCG only. † vs day 22.

**Table 4.** Differences in immune response to log-transformed ELISpot PPDA and PPDB for (i) BCG+GonaCon compared to BCG only and (ii) GonaCon only compared to BCG only during phase 1.

| Model term | PPDA |  |  | PPDB |  |  |
| --- | --- | --- | --- | --- | --- | --- |
|  | Coefficient | SE | P value | Coefficient | SE | P value |
| BCG+GonaCon* | -0.083 | 0.189 | 0.662 | -0.159 | 0.207 | 0.442 |
| GonaCon* | -0.813 | 0.218 | <0.001 | -1.118 | 0.238 | <0.001 |
| Day 42† | -0.283 | 0.179 | 0.114 | -0.206 | 0.201 | 0.305 |
| Day 63† | -0.583 | 0.179 | 0.001 | -0.639 | 0.201 | 0.001 |
| Day 90† | -0.526 | 0.179 | 0.003 | -0.457 | 0.201 | 0.023 |
| BCG+GonaCon: Day 42‡ | -0.173 | 0.226 | 0.939 | -0.153 | 0.254 | 0.546 |
| BCG+GonaCon: Day 63‡ | 0.203 | 0.226 | 0.369 | 0.245 | 0.254 | 0.336 |
| BCG+GonaCon: Day 90‡ | 0.144 | 0.226 | 0.526 | 0.153 | 0.254 | 0.546 |
| GonaCon: Day 42‡ | 0.873 | 0.261 | 0.001 | 0.955 | 0.293 | 0.001 |
| GonaCon: Day 63‡ | 0.488 | 0.261 | 0.061 | 0.582 | 0.293 | 0.047 |
| GonaCon: Day 90‡ | 0.644 | 0.261 | 0.013 | 0.677 | 0.293 | 0.021 |

\* vs BCG only. † vs day 22. ‡ vs main effects of treatment and day only.

##### PPDA

The analysis found no significant difference in PPDA response between BCG only and BCG+GonaCon, when measured using both IGRA and ELISpot. The coefficient for vaccination with GonaCon only compared to BCG only was negative and highly significant for both IGRA PPDA (p=0.003) and ELISpot PPDA (p<0.001). However, when measured by ELISpot this large difference was only apparent at day 22 and was not observed at day 42 (Figure 1b), meaning that additional interaction terms were required to model the relationships between treatment and day (Table 4). Immune responses in the BCG only group compared to GonaCon only were significantly different at all three timepoints (day 42, 63 and 90) compared to day 22 for the IGRA; for ELISpot, the day 63 and 90 terms were significant.

##### PPDB

Patterns observed for immune responses to PPDB were similar to PPDA in all treatment groups, as measured by both IGRA and ELISpot, in terms of the magnitude, direction and statistical significance of the model coefficients (Tables 3 and 4).

#### 3.1.2. Treatment effects on GonaCon response

No anti-GnRH antibody titres were detected at day 0 (for all treatment groups) nor at any sampling point in the BCG only group. All individuals from both the GonaCon only and the BCG+GonaCon group exhibited anti-GnRH antibody titres of at least 1:256,000 at the first sampling point, 22 days after administration of the first treatment dose (Table 5). High titre levels were maintained throughout phase 1, with only one of the 15 badgers (6.7%) in the BCG+GonaCon group and none of the badgers in the GonaCon only group dropping below 1:128,000 by day 90.

**Table 5.** Anti-GnRH titre levels (1:X,000) detected after initial treatment dose (at day 22, 42, 63, 90) in the BCG+GonaCon and GonaCon only groups. No anti-GnRH antibody titres were detected in any of the BCG only badgers and are therefore not included.

| Treatment | Badger ID | Sex | Day 22 | Day 42 | Day 63 | Day 90 |
| --- | --- | --- | --- | --- | --- | --- |
| BCG+GonaCon | 106 | F | 4096 | 8192 | 4096 | 2048 |
|  | 603 | F | 4096 | 2048 | 1024 | 512 |
|  | 96 | F | 1024 | 512 | 512 | 512 |
|  | 330 | F | 2048 | 1024 | 512 | 512 |
|  | 513 | F | 1024 | 1024 | 1024 | 512 |
|  | 625 | F | 1024 | 512 | 512 | 512 |
|  | 636 | F | 1024 | 512 | 256 | 128 |
|  | 778 | F | 512 | 512 | 512 | 512 |
|  | 540 | F | 512 | 256 | 128 | 64 |
|  | 53 | M | 2048 | 1024 | 512 | 512 |
|  | 256 | M | 1024 | 512 | 512 | 256 |
|  | 534 | M | 512 | 512 | 256 | 256 |
|  | 822 | M | 512 | 256 | 256 | 256 |
|  | 870 | M | 1024 | 1024 | 1024 | 1024 |
|  | 549 | M | 256 | 256 | 256 | 128 |
| GonaCon only | 377 | F | 1024 | 512 | 256 | 256 |
|  | 785 | F | 1024 | 512 | 256 | 128 |
|  | 807 | F | 512 | 256 | 128 | 128 |
|  | 852 | F | 512 | 256 | 256 | 256 |
|  | 556 | M | 2048 | 1024 | 1024 | 512 |
|  | 881 | M | 2048 | 4096 | 1024 | 1024 |
|  | 615 | M | 1024 | 1024 | 1024 | 1024 |
|  | 592 | M | 512 | 512 | 256 | 256 |

Average anti-GnRH antibody titres were plotted for the treatment groups (Figure 4); both GonaCon treatment groups produced a similar profile, peaking at the first sampling timepoint and declining gradually thereafter. The average anti-GnRH antibody titres for the BCG+GonaCon treated badgers were slightly higher than the GonaCon only treated badgers over the course of phase 1 (Table 6). The AIC for models 1, 2 and 3 were 214.3, 212.6 and 219.2 respectively. The higher AIC for model 3 suggested that the time effect was the same for both GonaCon and BCG+GonaCon groups. There was no clear preference between models 1 and 2, however they both provided similar conclusions about the effect of treatment on titre. Model 1 was chosen because long term extrapolation of the linear time effect in model 2 may give unreliable results. Regression coefficients from model 1 are presented in Table 7, which showed that the difference in titre between treatments was not statistically significant.

**Table 6.** Log_2_ transformed anti-GnRH antibody titres detected after vaccination with BCG+GonaCon and GonaCon only at days 22, 42, 63, 90. Values are means with standard deviations in parentheses.

| Days after vaccination | BCG+GonaCon | GonaCon only |
| --- | --- | --- |
| 22 | 10.000 (1.134) | 9.875 (0.834) |
| 42 | 9.467 (1.302) | 9.375 (1.302) |
| 63 | 9.000 (1.195) | 8.625 (1.188) |
| 90 | 8.533 (1.246) | 8.375 (1.188) |

**Table 7.** Differences in average log_2_ transformed anti-GnRH antibody titres between BCG+GonaCon and GonaCon only treatments during phase 1.

| Model Term | Coefficient | SE | P value |
| --- | --- | --- | --- |
| BCG+GonaCon vs. GonaCon only | 0.188 | 0.463 | 0.685 |
| Day 42<sup>†</sup> | -0.522 | 0.148 | <0.001 |
| Day 63<sup>†</sup> | -1.087 | 0.148 | <0.001 |
| Day 90<sup>†</sup> | -1.478 | 0.148 | <0.001 |

<sup>†</sup> vs day 22

**Figure 4.**
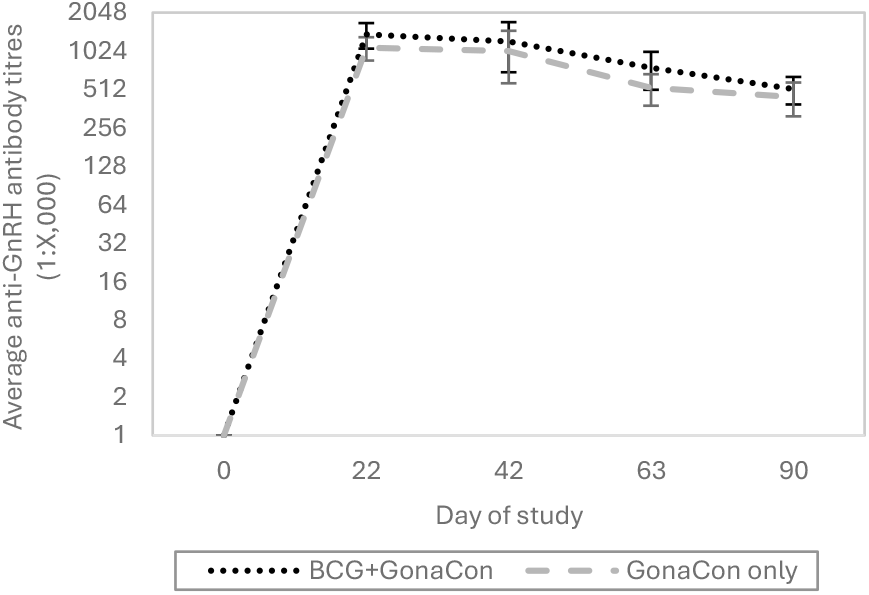
Average anti-GnRH antibody titres produced over time for the BCG+GonaCon and GonaCon only groups. No anti-GnRH antibody titres were detected in any of the BCG only badgers and are therefore not included. Error bars are standard error of the means.

### 3.2. Phase 2

#### 3.2.1. Immune response to BCG vaccination over time

The PPDA and PPDB immune responses to BCG vaccination over time (following initial BCG dose at day 0 and booster dose at day 373) for the six badgers from the BCG only treatment group are shown in Figures 5 and 6, respectively. Immune responses to both PPDA and PPDB were elevated at the first timepoint after initial vaccination (22 days) and booster vaccination (26 days), generally declining over time, although some individual profiles fluctuated during phase 1. The longer gap between blood sampling during phase 2 (26 days after BCG booster to day 271) provided less information for a comparable trend. There was a general consistency in the individual profiles as measured by both IGRA and ELISpot (Figure 7) immediately after vaccination, with all six badgers showing a peak in PPDB after initial and booster dosing; five of the six badgers also showed a similar pattern in PPDA response (animal ID803 being the exception). Immune responses to both PPDA and PPDB remained above pre-vaccination levels (day 0, phase 1) when measured 181 days after initial vaccination in five of the six badgers (ID350 was elevated for PPDA only), when measured using ELISpot. However, immune responses in the same five badgers had mostly returned to around pre-vaccination levels or below by 373 days after initial vaccination. By comparison, immune responses to PPDA and PPDB remained above pre-booster vaccination levels (day 0, phase 2) when measured 271 and 368 days after booster vaccination in five of the six badgers (ID259 being the exception), when measured using ELISpot.

**Figure 5.**
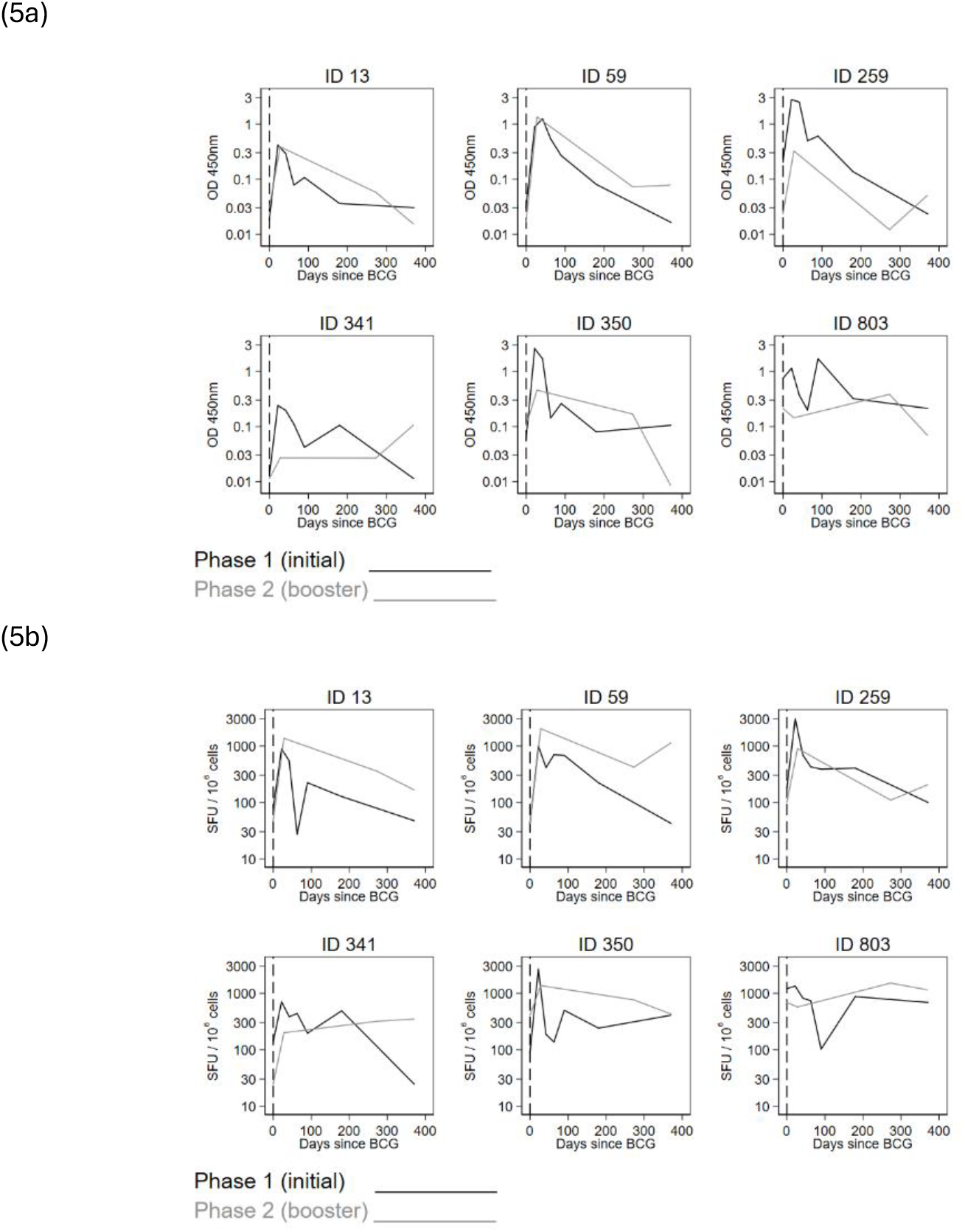
Individual profiles after initial and booster BCG vaccinations for (a) PPDA measured by IGRA and (b) PPDA measured by ELISpot.

**Figure 6.**
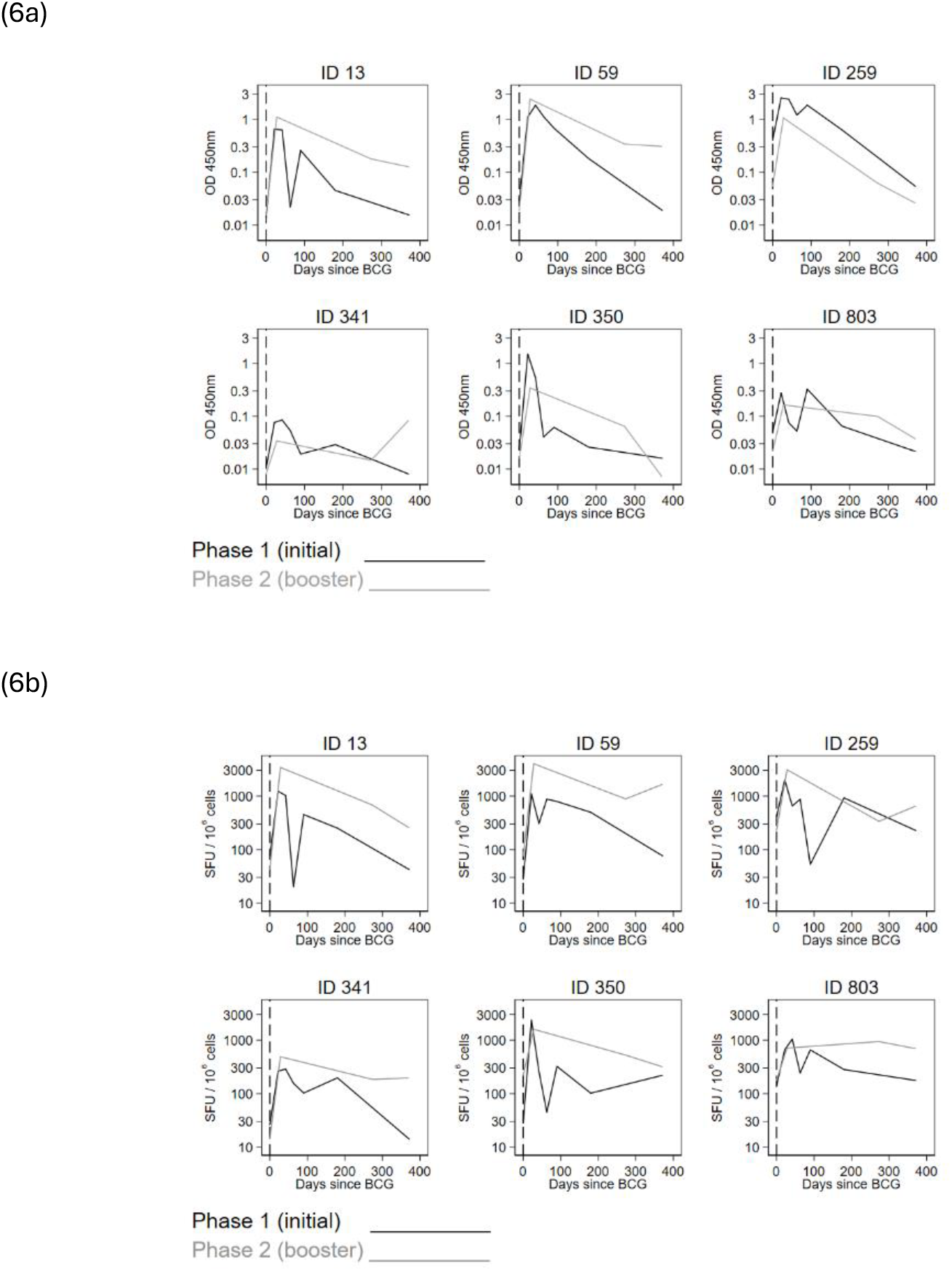
Individual profiles after initial and booster BCG vaccinations for (a) PPDB measured by IGRA and (b) PPDB measured by ELISpot.

**Figure 7.**
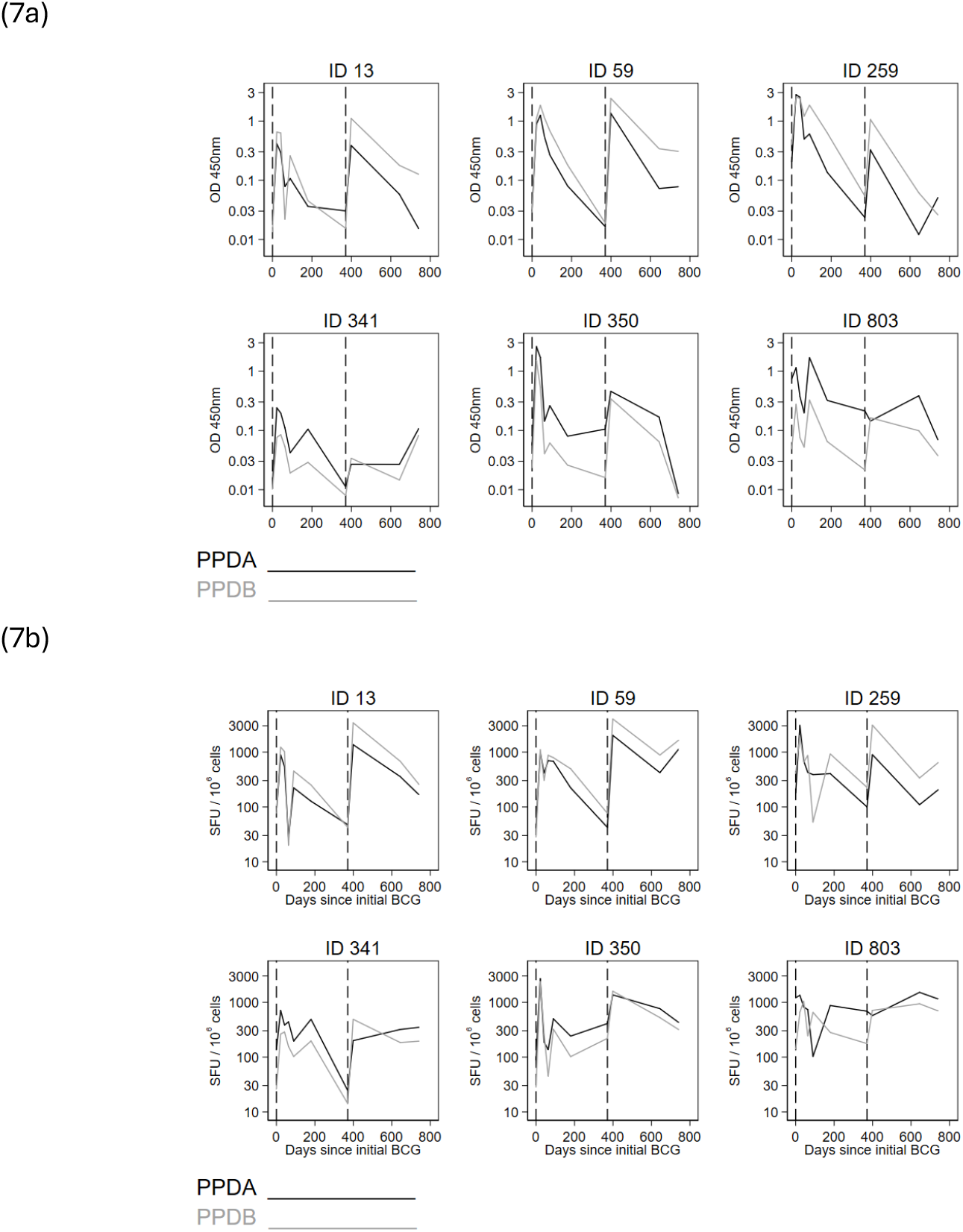
Individual (a). IGRA PPDA and PPDB and (b). ELISpot PPDA and PPDB profiles for the six badgers in BCG only group. Vertical lines denote BCG vaccination at day 0 (initial vaccination) and day 373 (booster vaccination).

## 4. Discussion

The pilot studies presented here aimed to determine the potential interactions and effect on immune responses when BCG and GonaCon were administered simultaneously in captive badgers. There were no notable deleterious effects on either vaccine, as the immune responses induced by both BCG and GonaCon did not differ statistically depending on whether they were administered on their own or together. The immune response to PPDB, which is considered a relatively strong indicator of effective vaccination against bTB, was greatest in the BCG only treatment group and lowest in the GonaCon only group, except the ELISpot peak at 42 days. However, the addition of GonaCon alongside BCG vaccination induced an immune response similar to BCG only, suggesting that GonaCon did not affect the immune response to BCG vaccination when both vaccines were administered together. Likewise, the anti-GnRH antibody titres generated by GonaCon only and BCG+GonaCon treatments were similar throughout the duration of phase 1, with the majority of badgers in both groups maintaining high titres (of 1:128,000 or above by day 90) that are associated with infertility in badgers (Cowan et al., 2019). Where titre levels declined, it is possible that these individuals would have returned to a fertile state although it has been demonstrated in several species, including badgers (Cowan et al., 2019), that infertility or reduced fecundity can persist at titres below a posited threshold (e.g. captive wild boar: Massei et al., 2012; rats: Pinkham et al., 2022), more so in captivity than field trials (Massei et al., 2012). Although these preliminary results are encouraging, a clear limitation of this study is that tests based on the measurement of immune responses alone do not indicate the protective effect afforded by either vaccine, in terms of reduced disease burden or fertility and further challenge studies (both *M. bovis* and breeding) would be required to demonstrate vaccine efficacy.

Although the immune responses to PPDA and PPDB were lower for the GonaCon only group compared to BCG only and BCG+GonaCon, there was a notable peak for both antigens at 42 days when measured using ELISpot. The adjuvant component of the GonaCon vaccine, AdjuVac, is a mineral oil-based surfactant adjuvant that contains killed *M. avium* as an immunostimulant (Miller et al., 2013), hence its reactivity with the PPDA antigen; the IFNγ responses to PPDB are probably reflective of cross-reactivity with environmental mycobacterial antigens (Rhodes et al., 2014).

The second phase of the study investigated the duration of immune response to BCG over 12 months followed by a booster dose. The results suggested that the immune response to BCG vaccination lasted no more than 12 months but could be effectively boosted by revaccination. In all six badgers (from the BCG only treatment group), immune responses to PPDA and PPDB were elevated at the first timepoint with most remaining above pre-vaccination levels for six months but dropping to around or below pre-vaccination levels after 12 months. Following a booster dose, immune responses were elevated again at the first timepoint in all but one badger, where the PPDA response declined. A strong booster effect was evident in most of the badgers (using ELISpot) with immune responses to PPDB remaining above pre-booster vaccination levels at 12 months. Revaccination with BCG has been shown to enhance the protective effect against bTB in several species, including deer (Griffin et al., 1999), cattle (Buddle et al., 2003; Parlane et al., 2014) and possums (Corner et al., 2002), depending on the interval between vaccination. Southey and Gormley (1999) also reported an enhanced immune response to PPDB in badgers revaccinated with BCG at a four-month interval. These authors concluded that the PPDB/PPDA T cell stimulation index ratio increased following booster vaccination, suggesting that vaccination had enhanced and switched the responses of badgers from an *M. avium* type response to an *M. bovis* BCG type response. Annual vaccination of badgers with BadgerBCG is already recommended on a population basis in view of the 30% rate of turnover, including new cubs and badger movement (VMD, 2020) but it could also enhance the duration of immunity in previously vaccinated individuals.

Since BadgerBCG was licensed in 2010, a variety of non-governmental groups and large-scale government operations have undertaken badger vaccination in England, concentrating on the High Risk Areas in south-west England (Benton et al., 2020). The current vaccination scheme allows trapping to be undertaken between May and November (open season); trapping is not permitted between December and April (closed season) due to inclement weather and potentially capturing pregnant or lactating females with dependent young. Benton et al. (2020) reported that captures peak around June and July, declining later in the autumn. Administering both GonaCon and BCG before or around this peak in summer would be advantageous for a number of reasons: previous research showed that delivering GonaCon to captive badgers earlier in the year (June) had an immediate effect on fertility and inhibited subsequent cub production the following year, compared to vaccination later in the year (November), which had no effect (Cowan et al., 2019). These authors suggested a possible explanation for this may be due to badgers’ unusual reproductive physiology and ability to delay implantation, whereby female badgers maintain fertilised eggs and dormant blastocysts at any time of the year, implanting around mid-winter (December to January) (Canivenc & Bonnin, 1981; Woodroffe, 1995; Dugdale et al., 2003). Early vaccination may have inhibited blastocyst production or affected the maintenance of blastocysts resulting in the absence of litters the following February/March. Cowan et al. (2019) concluded that the optimal window in which to administer GonaCon to badgers could be as narrow as between June and August as vaccinating badgers in November may be too late into the reproductive cycle to induce immediate infertility.

Vaccination with BCG early in the open season would also ensure cubs are targeted soon after weaning (May to June onwards), minimising the time during which cubs could be exposed to potential sources of *M. bovis* infection above ground (Carter et al., 2018). However, prior to this, cubs remain underground for the first two months of their life (Neal & Cheeseman, 1996); as badgers cannot be vaccinated until May at the earliest, there is a period after birth when cubs may be particularly susceptible to infection from close contact with other group members. Carter et al. (2012) reported that vaccinating at least one third of a social group reduced the risk of infection to unvaccinated cubs by 79%. This indirect benefit of vaccination for susceptible individuals highlights the importance of revaccinating badgers at least annually to maintain “herd immunity”, thus reducing the risk of *M. bovis* infection in the following year’s cubs. Revaccination with GonaCon would also boost the contraceptive effect: several studies in other species (e.g. white-tailed deer: Walker et al., 2021; feral cattle: Massei et al., 2018; feral horses: Baker et al., 2023) have shown that a booster dose of GonaCon significantly increases the proportion of animals rendered infertile and the duration of infertility. In badgers, vaccination would need to be repeated at least every two years to maintain the levels of female infertility predicted to have demographic impacts on badger populations (Cowan et al., 2019). The results from the current study and previous research suggests that a combined BCG and GonaCon vaccination campaign is likely to be most effective during the summer months (June to August), when capture rates of badgers peak and the likelihood of achieving infertility in female badgers is optimal.

The study of BCG vaccine and booster longevity was limited by the small sample size of animals in phase 2 and the lack of data during the early part of phase 2. As the objective of phase 2 was to investigate the longevity of immune response to BCG vaccination, it was only possible to include badgers from phase 1 that had received BCG only vaccination. Due to unforeseen circumstances, three of the original nine badgers from this cohort could not be included in the study, further reducing the sample size. Likewise, due to Covid-19 restrictions during phase 2, it was not possible to replicate the blood sampling schedule of phase 1, which limited the amount of information about the trend in phase 2.

A number of studies have reported on the effects of co-administering different vaccines, for example, foot-and-mouth vaccine and anthrax vaccine in cattle (Trotta et al., 2015) and sheep (Çokçalişan, et al., 2019), GonaCon and rabies vaccine in feral dogs (Bender et al, 2009) and feral cats (Novak et al., 2021) with no adverse effects on the target vaccine observed in any of these studies. Bender et al. (2019) reported that the use of GonaCon did not affect the ability of dogs to seroconvert in response to the rabies vaccine, concluding that GonaCon used in combination with rabies vaccine had the potential to increase herd immunity and control overabundance of the dog population to enhance the management of rabies. Despite the limitations of the current study, the minimal interference seen between BCG and GonaCon when administered together, suggests that a similar scenario could be achieved with badgers to effectively control badger populations and manage bTB simultaneously. These preliminary results suggest that further captive or field investigations on a larger scale would be beneficial.

## 5. Acknowledgements

We are grateful to Richard Delahay (APHA, Woodchester Park, UK) and Mark Chambers (APHA, Weybridge, UK) for their comments on manuscript drafts. The study was carried out under a UK Home Office licence, in accordance with the Animals (Scientific Procedures) Act 1986 and was approved by the APHA’s Ethical Review Process PBDA2E164-4-001.

